# Lytic phages displayed protective effects against soft rot-causing *Pectobacterium* sp.

**DOI:** 10.1101/2021.11.15.468558

**Authors:** Aeron Jade S. Parena, Benji Brayan I. Silva, Rae Mark L. Mercado, Adelbert Adrian A. Sendon, Freddiewebb B. Signabon, Johnny F. Balidion, Jaymee R. Encabo

## Abstract

Soft rot caused by *Pectobacterium* sp. is responsible for significant losses in vegetable production worldwide. Methods for the effective control of soft rot are limited and are primarily based on good agricultural practices. The use of bacteriophages as biocontrol agents appears to be a promising alternative to combat phytopathogens. In this study, we investigated the efficacy of lytic phages recovered from symptomatic tissues and environmental samples against soft rot caused by *Pectobacterium carotovorum* subsp. *carotovorum*. Three bacteriophage isolates, designated as vB_PcaP-A3, vB_PcaM-D1, and vB_PcaM-J3, were observed to effectively lyse *P. carotovorum* subsp. *carotovorum*. Phage vB_PcaP-A3 exhibited virion morphology similar to the members of the podovirus group, while phages vB_PcaM-D1 and vB_PcaM-J3 showed myovirus morphology based on transmission electron microscopy. The optimal multiplicity of infection (MOI) differed greatly among the three phages. All three phages survived incubations at 30°C, 40°C and 50°C and pH conditions ranging from 3.0 to 9.0, but were all inactivated at 60°C and at pH 12. Both monophage and cocktail preparations were effective in inhibiting the growth of *P. carotovorum* subsp. *carotovorum* in the *in vitro* challenge tests. In the semi-*in planta* assays, while treatment with cocktail preparations completely inhibited the development of soft rot in tissue slices, monophage treatments not only resulted in significant reduction of tissue maceration in slices, but also showed protective effect against soft rot in tubers. Overall, these results demonstrate the efficacy of phages vB_PcaP-A3, vB_PcaM-D1, and vB_PcaM-J3 for the biocontrol of soft rot caused by *P. carotovorum* subsp. *carotovorum*.

## 1. Introduction

Bacterial soft rot disease causes serious damage among vegetable and fruit crops worldwide (Bhat et al., 2010). Annual post-harvest losses due to bacterial soft rot vary from 15-30% of the harvested crops and at least 30-50% of the 65 million tons of vegetables in public markets (Bhat et al., 2010; Monjil et al., 2021; Rahman et al., 2012). Soft rot disease results from infections by several types of bacteria, but most commonly by members of the soft rot *Pectobacteriaceae* (SRP) including *Pectobacterium carotovorum* subsp. *carotovorum, P. aroidearium, P. atrosepticum, P. brasiliense, P. betavasculorum* and *P. wasabiae*, which were identified to cause soft rot both in storage and in transit (Adeolu et al., 2016; Agyemang et al., 2020; Charkowski, 2018; Czajkowski, 2015; Gardan et al., 2003; Nabhan et al., 2013; Pérombelon, 2002; Portier et al., 2019; Wallace, 1948). *Pectobacterium* species have the broadest host range among soft rot bacteria, and are considered as the leading cause of soft rot disease in various vegetable crops (Davidsson et al., 2013; Lim et al., 2017). During infection, SRPs produce enzymes like pectate lyase, cellulase and polygalacturonase, which are capable of breaking down the plant cell wall leading to plant tissue maceration and decay (Charkowski et al., 2012; Li et al., 2019; Mattinen et al., 2007; Muturi et al., 2019). There is still limited information on the predominant SRPs affecting vegetable crops in the Philippines. A previous report by Daengsubha and Quimio (1980) identified *P. carotovorum* subsp. *carotovorum* as the major cause of bacterial soft rot affecting crucifers, carrot, and tomato.

Methods used to control soft rot disease include cutting-off and disposal of infected tissues during or after harvest, wrapping with paper, treatment with hot water, and the use of bactericides, such as alum powder (aluminum potassium sulfate), formaldehyde, copper sulfate, and sodium or calcium hypochlorite (Bayogan et al., 2011; Bhat et al., 2010; Czajkowski et al., 2011; Liao et al., 2003). However, these methods only slow down the development of soft rot, and the use of chemical bactericides were reported to be associated with the detection of copper- and antibiotic-resistant phytopathogenic bacterial strains (Behlau et al., 2011; Cooksey, 1990; McManus et al., 2002; Petriccione et al., 2017; Roach et al., 2020; Sundin & Wang, 2018). Due to the limited success of the abovementioned control methods, and the currently growing global threat of antimicrobial resistance, there has been an increased interest in the use of bacteriophages or phages (viruses of bacteria) as biocontrol agents against pathogenic bacteria (Buttimer et al., 2017; Frampton et al., 2012; Jones et al., 2007; Jones et al., 2012).

Bacteriophages (phages) are viruses that infect and lyse bacterial cells. When environmental conditions are favorable for infection, phages can effectively reduce bacterial populations, as in the case of lytic phages that undergo rapid replication (Hadas et al., 1997; Sulakvelidze et al., 2001). Since phages have high host-specificity and have no direct negative effects on non-host cells, these allow the use of phage therapy to be a promising biocontrol method against soft rot caused by *Pectobacterium* sp. (Buttimer et al., 2017; Lim et al., 2017). The first *Pectobacterium* phage that was sequenced, PP1, was reported in 2012 and several additional novel phage isolates were subsequently reported (Buttimer, Lucid, et al., 2018; Buttimer et al., 2020; Kalischuk et al., 2015; D. H. Lee et al., 2012; Lim et al., 2017; Lim et al., 2013; Lim et al., 2015; Lim et al., 2014; Pedersen et al., 2020; Voronina et al., 2019). These bacteriophages vary especially in terms of morphology, lytic activity and host range but they all have the potential to control soft rot caused by *Pectobacterium* sp. as evidenced by the semi-*in planta* experiments performed in these reports.

In this study, three lytic phages isolated from symptomatic tissues and environmental samples, were characterized based on virion morphology, growth characteristics, and physico-chemical properties. The lytic activity and efficacy of the phages to prevent tissue maceration caused by *P. carotovorum* subsp. *carotovorum* in potato slices using monophage and cocktail suspensions were also determined. The effect of the timing of phage application on the efficacy of monophage treatments on whole potato tubers was also qualitatively assessed. Based on the *in vitro* and semi-*in planta* assays, we report the biocontrol potential of these lytic phages against bacterial soft rot caused by *P. carotovorum* subsp. *carotovorum*.

## 2. Materials and Methods

### 2.1. Bacterial Strains and Media

*Pectobacterium carotovorum* subsp. *carotovorum* BIOTECH 1752, obtained from the Philippine National Collection of Microorganisms (National Institute of Molecular Biology and Biotechnology, UPLB) was used as the primary host for phage isolation. A collection of 48 *Pectobacterium* spp. isolated from vegetable growing areas and market centers in the Philippines (Host-Pathogen Interaction Laboratory, IWEP, UPLB), and eight other representative strains from other genera (Microbiology Division, IBS, UPLB) were used for the host range assay (Supplementary Table 1). For routine tests and maintenance, bacteria were grown in Tryptone Soya Agar (TSA, HiMedia®, Mumbai, India) for 16-18 h at 30°C. For liquid preparations, bacterial cultures were grown in Tryptone Soya Broth (TSB, HiMedia®, Mumbai, India) for 12-18 h at 30°C with 125 rpm agitation.

### 2.2. Bacteriophage Isolation and Characterization

#### 2.2.1. Isolation and Purification of Bacteriophages

Bacteriophages were isolated from rhizosphere soil, symptomatic tissues, and sewage water (Czajkowski et al., 2014). Twenty grams (or mL) of each sample was homogenized in 20 mL saline-magnesium (SM) buffer (MgSO_4_, NaCl, CaCl_2_, and Tris-HCl, pH 7.5) and incubated at 30°C for 12-16 h. Five hundred μL of log phase (5 to 6-h subculture prepared from an overnight culture) *P. carotovorum* subsp. *carotovorum* culture containing approximately 10^8^ cells mL^-1^ (0.5 McFarland Standard) was added to each enrichment set-up and was incubated as above. Enrichment cultures were centrifuged at 3,000 rpm for 15 min and the supernatants were chloroform-treated in 1:20 chloroform/supernatant ratio for at least 10 min. The resulting lysates were screened for the presence of bacteriophages by spot test assay (Mazzocco et al., 2009). Overlay mixtures, containing 4.5 mL soft agar (TSA), 0.5 mL 20 mM CaCl_2_, and 300 µL of log phase *P. carotovorum* subsp. *carotovorum* culture (0.5 MFS) were poured on Petri plates containing pre-solidified TSA. Five μL of the phage lysates were spotted on the overlay and the spots were absorbed into the agar for 10-15 min prior to incubation of plates at 30°C for 12-16 h. Zones of lysis on the bacterial lawn indicated the presence of phages.

Phage purification was done using the double layer plaque assay (Kropinski et al., 2009). Ten-fold dilutions were prepared from each positive lysate. In separate sterile 1.5-mL microcentrifuge tubes, 300 μL of diluted lysate suspensions were mixed with 300 μL log phase culture of *P. carotovorum* subsp. *carotovorum* (0.5 MFS) in TSB. These were pre-incubated for 10 min at 4°C to allow adsorption of the phages. Molten overlay (4.5 mL) mixed with 0.5 mL 20 mM CaCl_2_ was added to the suspensions and the resulting mixtures were poured onto Petri plates containing pre-solidified TSA. The overlays were allowed to solidify for 15-20 min and the plates were incubated at 30°C for 12-16 h. Well-isolated plaques were picked using sterile pipette tips and the phages were suspended in 500 µL SM buffer. Bacteriophage isolates were purified by successive single plaque isolation for at least five times or until uniform plaque morphology was observed.

#### 2.2.2. Plaque Formation Using Double Layer Plaque Assay

Plaque morphologies were characterized after a series of double layer plaque assays as described above (Jurczak-Kurek et al., 2016). The transparency and morphology of the plaques formed by each bacteriophage isolate was described and plaque diameter was measured using a digital caliper.

#### 2.2.3. Bacteriophage Lysate Preparation and Titration

Double layer plaque assay plates with uniform plaques were flooded with 3-5 mL of SM buffer to obtain the bacteriophage lysate. Plates were gently shaken for 15-20 min and the overlay-SM buffer mixtures were collected and centrifuged at 8,000 rpm for 10 min. The supernatants were chloroform treated in 1:20 chloroform/lysate ratio and were maintained at 4°C until titration. Lysates were subjected to double layer plaque assay (or spot test) for titration as described above. Phage titer was determined and expressed as plaque forming units (PFU mL^-1^).

#### 2.2.4. Determination of Virion Morphology Using Transmission Electron Microscopy (TEM)

High titer lysates (at least 10^10^ PFU mL^-1^) of the three phage isolates were submitted to the Electron Microscopy Laboratory of the Research Institute for Tropical Medicine (Muntinlupa City, Philippines). The phage suspension was placed on carbon-coated copper grids, and 2% aqueous uranyl acetate was added for three min to negatively stain the virions. Phages were visualized by transmission electron microscopy (TEM; JEOLJEM-1220 Electron Microscope), and the images were scanned with 800 ms exposure, 1.4 gain, and 1.0 bin with no sharpening, normal contrast, and 1.00 gamma.

#### 2.2.5. Optimal Multiplicity of Infection (MOI)

The optimal ratio of phage particles to the number of bacterial cells, also called the optimal multiplicity of infection (MOI) was determined for the three phage isolates. *P. carotovorum* subsp. *carotovorum* was infected with each phage isolate at four different PFU mL^-1^/cell mL^-1^ ratios (MOI): 0.01, 0.1, 1.0, and 10. Following overnight incubation at 30°C, the suspensions were chloroform-treated (1:20) for 10 min and centrifuged at 8,000 rpm for 10 min. The supernatants were subjected to double layer plaque assay to determine phage titer. The MOI demonstrating the highest phage titer or PFU mL^-1^ was considered optimal.

#### 2.2.6. Latent Period and Phage Burst Size

One-step growth experiments were conducted to determine phage latent period and burst size following previously described protocol (Czajkowski et al., 2014). Cells of *P. carotovorum* subsp. *carotovorum* at log phase were harvested by centrifugation at 12,000 rpm for 15 min at 10°C, and were resuspended in TSB to an OD_600_ of 0.5 (c. 5 × 10^8^ CFU mL^-1^). Subsequently, 100 μL of 20mM CaCl_2_ and 100 μL of the phage isolate were added into the bacterial culture at MOI of 0.1 (10^7^ PFU mL^-1^/ 10^8^ CFU mL^-1^). The phages were allowed to adsorb by incubation at 4°C for 10 min and the suspensions were further incubated at 30°C for 10 min. The suspensions were centrifuged at 12,000 rpm for 15 min at 10°C to remove unadsorbed phages. The resulting pellet was resuspended in 10 mL fresh TSB. The mixtures were incubated at 30°C for 60 min with agitation at 125 rpm. Aliquots were taken from the mixtures every 10 min and the titer was determined using the Spot Test method. The burst size was calculated using the formula: 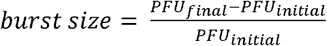 and the experiment was repeated twice with the same set-up and the results were averaged.

#### 2.2.7. Adsorption Rate

Phage adsorption rate was determined following previously described protocol (Czajkowski et al., 2014). One mL of *P. carotovorum* subsp. *carotovorum* (ca. 10^8^ CFU mL^-1^) was infected with each phage isolate by adding 100 μL of phage lysate (ca. 10^8^ pfu mL^-1^) to reach MOI 0.1 and the resulting suspensions were incubated at 30°C for 20 min. After 0 (control), 1, 2, 4, 10 and 20 min, 50-μL aliquots were taken from each suspension and added to 450 μL TSB. Ten-fold dilutions in SM buffer were prepared and the residual titer of the unadsorbed phages were determined using the spot test method as described above. The experiment was repeated twice with the same set-up and the results were averaged. Phage adsorption was calculated as follows: 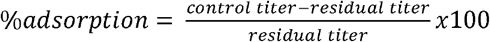

#### 2.2.8. Thermal Resistance and pH Stability

Thermal resistance of the three phages was determined at 30, 40, 50 and 60 °C. Phages (10^8^ PFU mL^-1^) in SM buffer were incubated for 1 h at each temperature condition and the surviving PFU was determined using double layer plaque assay (Czajkowski et al., 2014).

The effect of pH on phage stability was determined using phages (10^8^ PFU mL^-1^) in SM buffer within a pH range of 3.0 to 12.0. Phage lysates (10^8^ PFU mL^-1^) were added to SM buffer, adjusted with 2N HCl or 3N NaOH to pH 3, 7, 9, and 12 (Lim et al., 2013). The experiment was carried out by incubating the phage suspensions at 4°C for 16-24 h and the surviving PFU was determined using double layer plaque assay.

#### 2.2.9. Host Range Determination

The host ranges of the three phage isolates was determined using a collection of 48 *Pectobacterium* spp. isolates and eight representative strains from different genera (Supplementary Table 1) using the spot test assay. *Pectobacterium* spp. were isolated in the years 2017-2019 from symptomatic tissues of various vegetable crops.

For the spot test assay, bacterial cultures were grown in TSB at 30°C for 12-16 h with agitation at 125 rpm. From the resulting bacterial cultures, 300 μL (0.5 MFS) of each was inoculated into the overlay agar and was subsequently poured into a prepared TSA plate and allowed to solidify at room temperature for 30 min. Five μL of the phage lysate (10^9^ PFU mL^-1^) was spotted onto the top agar layer, and the plates were allowed to dry at room temperature for 10-15 min. Following overnight incubation at 30°C, plates were inspected for lysis zones.

### 2.3. In Vitro Challenge Test

Bacterial inactivation using monophage (vB_PcaP-A3, vB_PcaM-D1, or vB_PcaM-J3) and cocktail treatments (vB_PcaP-A3+D1, vB_PcaP-A3+J3, vB_PcaM-D1+J3, or vB_PcaP-A3+D1+J3) was determined following the method described by Chen et al. (2018). For monophage treatments, log phase culture of *P. carotovorum* subsp. *carotovorum* (OD_600_ of 0.5, c. 5 × 10^8^ CFU mL^-1^) was mixed with the phage lysate at optimal MOI. For cocktail preparations, low (0.001) and high (100) MOI were used for the experiment. The bacteria-phage mixtures were incubated at 30°C with agitation at 125 rpm and the OD readings at 600nm were recorded (SPECTROstar^*Nano*^, Ortenberg, Germany) at 15-minute intervals for eight hours to monitor the changes in bacterial density.

### 2.4. Semi-In Planta Assays

#### 2.4.1. Phage Biocontrol Using Monophage and Cocktail Treatments on Tissue Slices

The efficacies of monophage and cocktail treatments against soft rot caused by *P. carotovorum* subsp. *carotovorum* were evaluated using a potato slice assay (Czajkowski et al., 2015). Potato tubers (cultivar Igorota) were surface sterilized by soaking in 70% ethanol for 10 min, washed with sterile distilled water, and air dried under the laminar flow hood. The tubers were cut into 7.0 mm-thick transverse disks using sterile knives and three wells (5-mm diameter) were made on each slice using a sterile cork borer. For monophage treatments, log phase culture of *P. carotovorum subsp. carotovorum* containing approximately 10^8^ cells mL^-1^ (0.5 MFS) was mixed with the phage lysates at optimal MOI. For cocktail preparations, low (0.001) and high (100) MOIs were used for the experiment. Three potato slices were used for each treatment. Wells were filled with 20 µL of the bacteria-phage mixtures, 20 µL of sterile SM buffer as negative control, and 20 µL of *P. carotovorum* subsp. *carotovorum* (0.5 MFS) as positive control. The diameter of the rotten tissues around the wells inoculated with bacteria and wells co-inoculated with bacteria and phages were measured after incubation at 30°C for 72 h. Treatment means were compared using one-way ANOVA with Tukey’s post-test (α = 0.05).

#### 2.4.2. Protective and Curative Potential of Monophage Treatments against Soft Rot

A potato tuber assay was conducted to determine the protective and curative effect of monophage treatments at various application time points relative to *P. carotovorum* subsp. *carotovorum* inoculation following a previously described protocol by Czajkowski et. al. (2014). Potato tubers (cultivar Igorota) were washed in running water to remove soil and other organic particles, surface-sterilized with 70% ethanol for 10 min, rinsed with sterile water for three times, and air dried under laminar flowhood. For each tuber, two wells (8 mm diameter) were made using flame-sterilized cork borer. Two set-ups were prepared for the experiment: (1) advanced phage introduction (API) treatment, and (2) delayed phage introduction (DPI) treatment. For both treatments, log phase culture of *P. carotovorum subsp. carotovorum* containing approximately 10^7^ cells mL^-1^ was used and the phage lysate was added to the wells at MOI 10. In the API treatment, wells were filled with 25 μL of the phage suspension at 0, 1, 3, 8, 15 and 30 h prior to the inoculation of 25 μL of the bacterial suspension. In the DPI treatment, wells were filled with 25 μL of the phage suspension at 0, 1, 3, 8, 15 and 30 h after bacterial inoculation. Three potato tubers were used per time point for each of the API and DPI treatment. For the control set-ups, 25 μL aliquots of sterile water was used as negative control while 25 μL of *P. carotovorum* subsp. *carotovorum* suspension was used as positive control. The tubers were placed in a humid box and the degree of tissue maceration was qualitatively assessed (scored as +/-) after incubation at 30°C for 72 h.

## 3. Results

### 3.1. Isolation and Characterization of Bacteriophages

#### 3.1.1. Isolation of Lytic Bacteriophages

Bacteriophages vB_PcaP-A3, vB_PcaM-D1 and vB_PcaM-J3 were isolated from rhizosphere soil, symptomatic tissues, and sewage water, respectively, using *P. carotovorum* subsp. *carotovorum* as enrichment and isolation host. The three phages were named following the unified phage naming system proposed by Kropinski et al. (2009). All three phages formed clear plaques typical for lytic phages, with varying sizes (Table 1).

**Table 1.** Characteristics and biological properties of the three lytic bacteriophages isolated using *Pectobacterium carotovorum* subsp. *carotovorum*.

| Phage | Morphology | Head diameter<br>(nm) | Tail Length (nm) | Plaque Morphology |  | Optimal MOI | Latent Period (min) | Burst Size (PFU/cell) |
| --- | --- | --- | --- | --- | --- | --- | --- | --- |
|  |  |  |  | Transparency | Diameter (mm) |  |  |  |
| vB_PcaP-A3 | Podovirus | 81.07 ± 7.04 | 19.58 ± 2.92 | clear | 2.0 - 3.0 | 0.01 | 10 | 142 |
| vB_PcaM-D1 | Myovirus | 77.50 ± 4.68 | 98.15 ± 4.09 | clear | 1.0 - 1.9 | 0.1 | 10 | 151 |
| vB_PcaM-J3 | Myovirus | 76.17 ± 2.81 | 137.08 ± 12.78 | clear | < 1.0 | 10 | 30 | 42 |

#### 3.1.2. Virion Morphology

Examination of the phages using TEM revealed that vB_PcaP-A3 exhibited podovirus morphology while vB_PcaM-D1 and vB_PcaM-J3 showed virion morphology similar to members of the myovirus group (Table 1, Figure 1). Phage vB_PcaP-A3 has an icosahedral head of 80 nm and short non-contractile tail of 20 nm (Table 1). Phages vB_PcaM-D1 and vB_PcaM-J3 both have icosahedral heads of around 75 nm, with longer, rigid contractile tails that are about 95-135 nm (Table 1, Figure 1).

**Figure 1.**
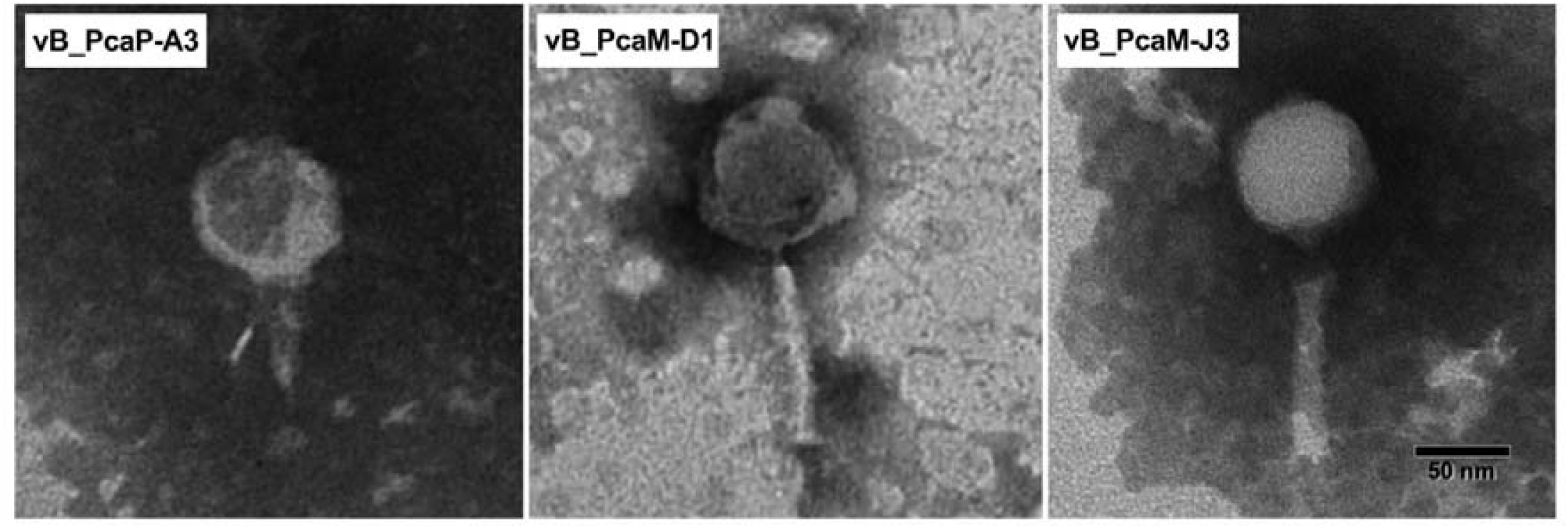
Transmission electron micrographs (TEM) of vB_PcaP-A3, vB_PcaM-D1, and vB_PcaM-J3 virions. High titer phage lysates were negatively stained with 2% aqueous uranyl acetate and were visualized by transmission electron microscopy (TEM; JEOL JEM-1220 Electron Microscope). The images were scanned with 800 ms exposure, 1.4 gain, and 1.0 bin with no sharpening, normal contrast, and 1.00 gamma. Bar represents 50nm.

#### 3.1.3. Characterization of Bacteriophages

All three phages exhibited strong adsorption to *P. carotovorum* subsp. *carotovorum*, and approximately 90% of the phage particles adsorbed to the bacterial cells within five min (Figure 2). vB_PcaP-A3 and vB_PcaM-D1 showed similar infection kinetics with a latent period of 10 min and a burst size of 142-151 progeny phages/cell, whereas vB_PcaM-J3 showed a longer latent period of 20 min and a smaller burst size of 42 progeny phages/cell (Table 1, Figure 2A & 2B). Among the three phages, vB_PcaP-A3 has the lowest optimal MOI (0.01), followed by vB_PcaM-D1 (1.0) and vB_PcaM-J3 (10.0) (Table 1). All of the three phages retained infectivity after incubation at 30°C, 40°C, and 50°C for 1 h, and in the pH range 3-9 for 16-24 h (Figure 2C).

**Figure 2.**
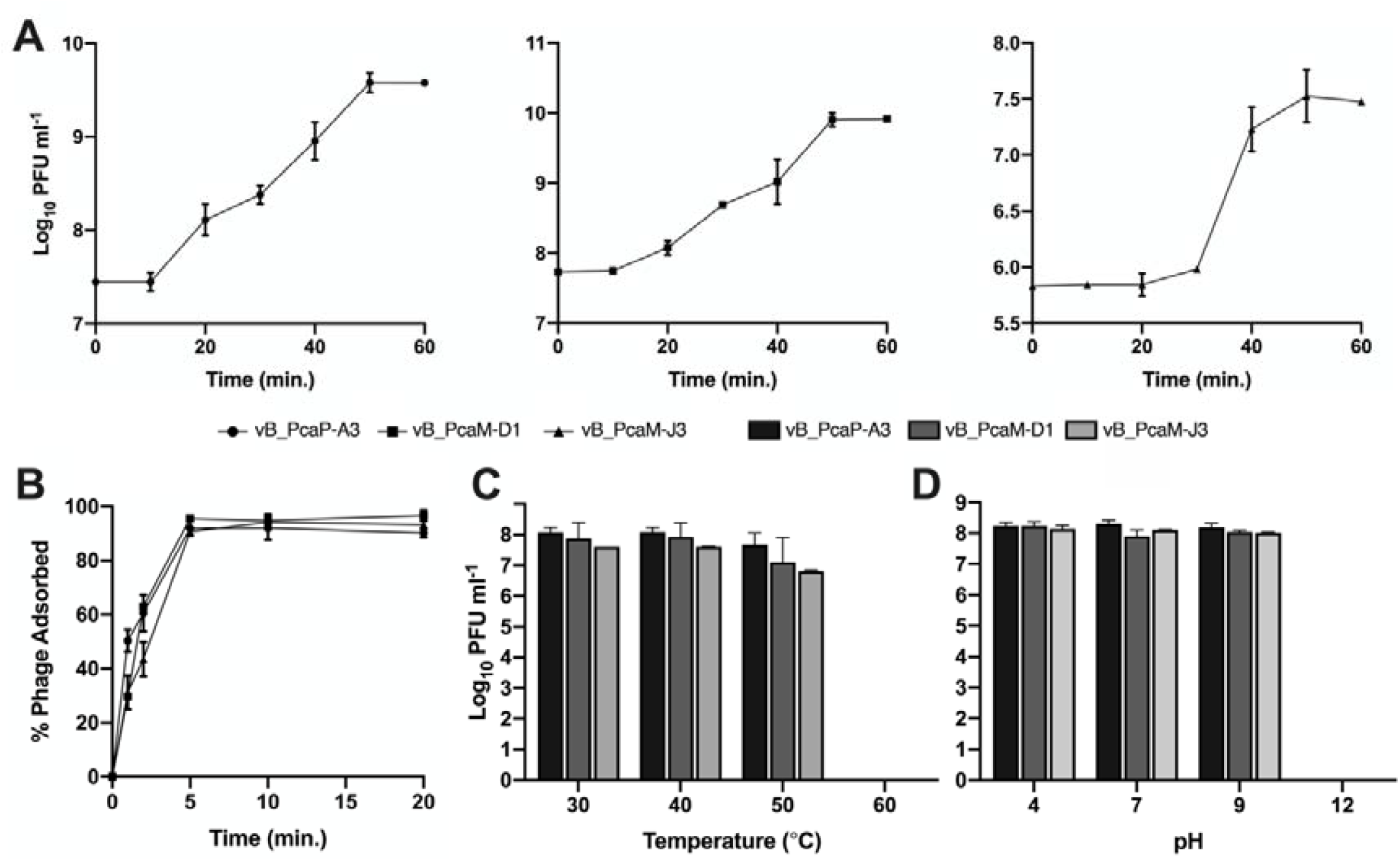
Growth characteristics of phages vB_PcaP-A3, vB_PcaM-D1, and vB_PcaM-J3 and their stability on varying temperature and pH levels. **(A)** One-step growth curve of each phage grown together with log phase *Pectobacterium carotovorum* subsp. *carotovorum* in TSB for 60 min at 30 °C. **(B)** Adsorption of each phage to log phase *Pectobacterium carotovorum* subsp. *carotovorum* in TSB for 60 min at 30 °C. **(C)** Stability of the phages at various temperature and pH levels.

#### 3.1.4. Host Range

Spot test assay revealed that the three phages showed antibacterial activity specific for *P. carotovorum* subsp. *carotovorum*. None of the three phages was able to lyse the 48 *Pectobacterium* spp. isolates and the eight representative strains from other genera.

#### 3.1.5. Phage Activity In Vitro

Killing curves established for vB_PcaP-A3, vB_PcaM-D1 and vB_PcaM-J3 showed that all three phages have strong lytic activity and were highly effective against *P. carotovorum* subsp. *carotovorum* at optimal MOI (Figure 3A). The bacterial cell density remained less than 0.05 (OD_600_) throughout the 8-h observation period. The absence of any increase in OD_600_ reading also suggests that there is no observed phage resistance development of the host, at least during the observation period (Muturi et al., 2019). These results indicate that monophage treatments of vB_PcaP-A3, vB_PcaM-D1 and vB_PcaM-J3 at optimal MOI were equally effective in inhibiting the growth of *P. carotovorum* subsp. *carotovorum in vitro*.

**Figure 3.**
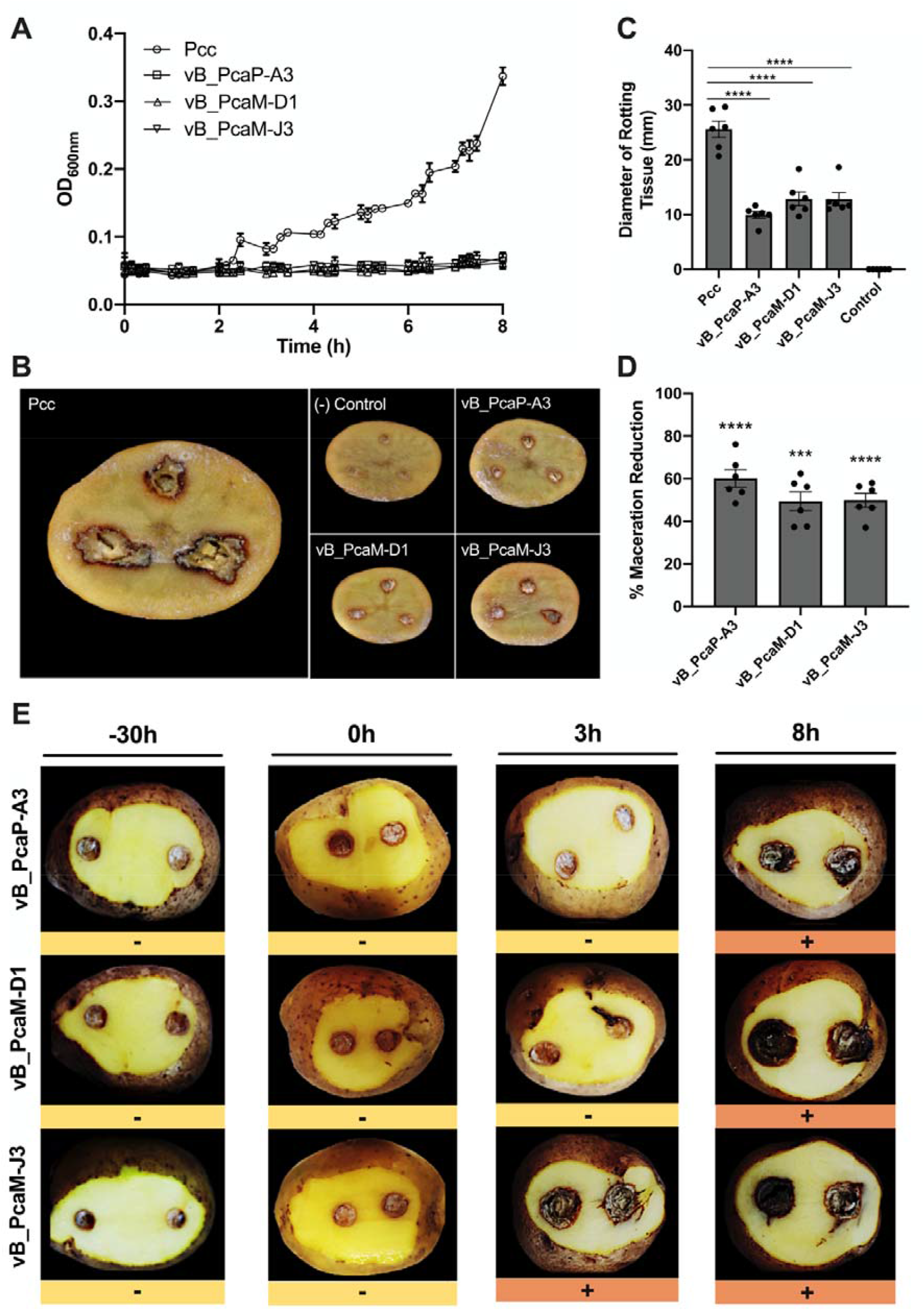
Bacterial killing curves of phages vB_PcaP-A3, vB_PcaM-D1, and vB_PcaM-J3 and their protective effect against *Pectobacterium carotovorum* subsp. *carotovorum* on potato tuber slices based on optimal MOI. **(A)** Killing curves of phages vB_PcaP-A3, vB_PcaM-D1, vB_PcaM-J3 against *Pectobacterium carotovorum* subsp. *carotovorum* measured as change in absorbance at OD_600 nm_. **(B)** Protective effect of monophage treatment on potato tuber slices against *Pectobacterium carotovorum* subsp. *carotovorum*. **(C)** Mean diameter of rotting tissue. **(D)** Percent tissue maceration reduction in potato slices co-inoculated with *Pectobacterium carotovorum* subsp. *carotovorum* and vB_PcaP-A3, vB_PcaM-D1 or vB_PcaM-J3 and **(E)** the preventive-curative effect of selected bacteriophages in potato tubers against *Pectobacterium carotovorum* subs. *carotovorum* after 72-h incubation at 28°C. Each dot in **(C-D)** is an individual potato slice (ie. replicate). Statistical significance was determined with one-way ANOVA with Tukey’s post-test and one-sample t-test in **(C)** and **(D)**, respectively, ****p < 0.0001.

For the cocktail treatments, low MOI (0.001) and high MOI (100) were used in the *in vitro* challenge tests. Each phage was used at the same concentration and the phage cocktails consisting of two or three phages were combined and evaluated. At MOI of 0.001, the killing curves of the phage cocktails composed of two or three phages maintained its bacterial inactivation with an OD_600_ < 0.05 for 8 h (Figure 4A). At low MOI, it appears that a phage cocktail containing any of the three phages effectively controlled bacterial growth during the 8-h observation period. At MOI 100, killing curves showed that all cocktail treatments were highly effective against *P. carotovorum* subsp. *carotovorum* (Figure 4A). The bacterial density remained less than 0.05 (OD_600_) throughout the duration of the experiment.

**Figure 4.**
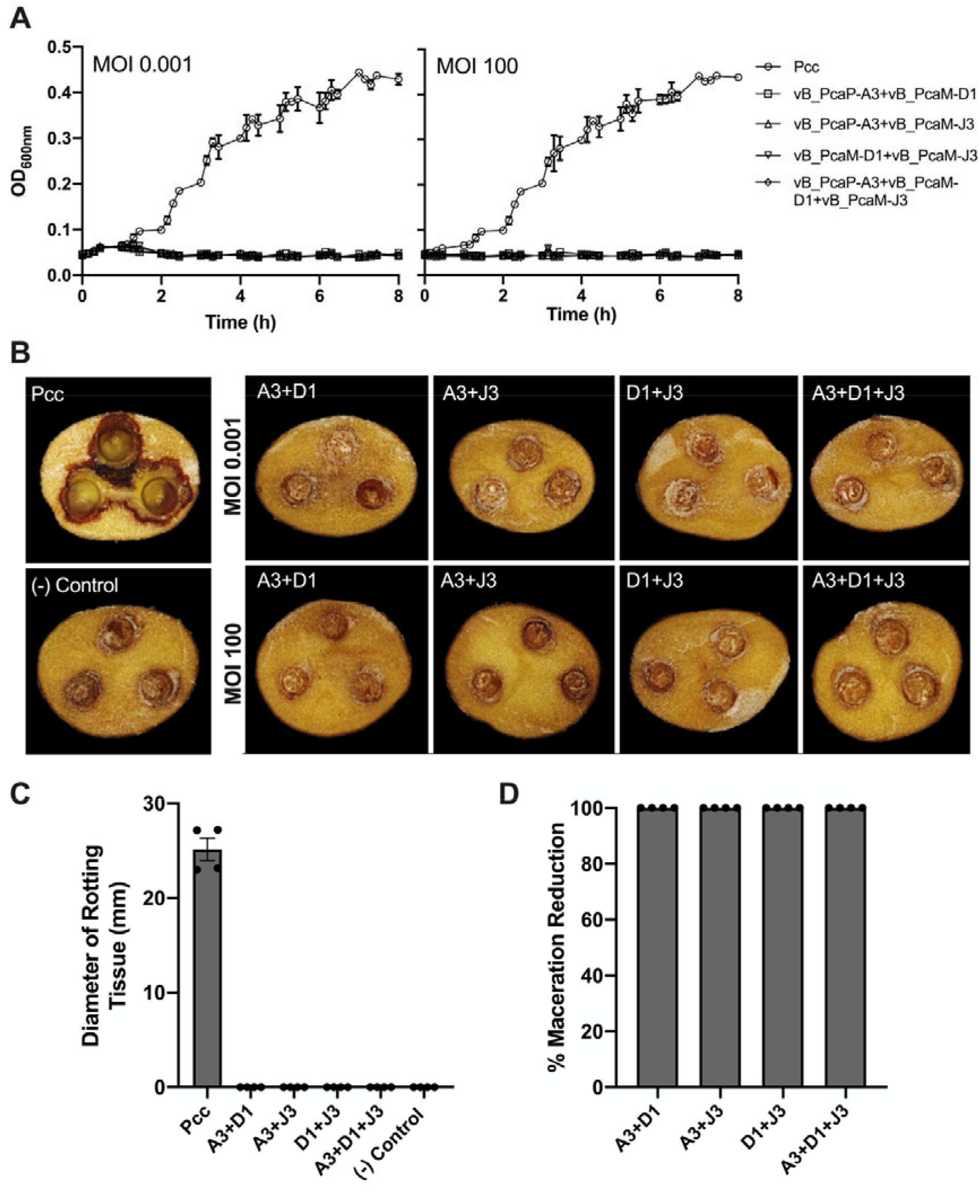
Suppression of *Pectobecterium carotovorum* subsp. *carotovorum* by phage cocktails *in vitro* and semi-*in planta*. **(A)** Lytic activities of cocktails of two (vB_PcaP-A3+vB_PcaM-D1, vB_PcaP-A3+vB_PcaM-J3, and vB_PcaM-D1+vB_PcaM-J3) and three (vB_PcaP-A3+vB_PcaM-D1+vB_PcaM-J3) phages at low and high MOI against *Pectobacterium carotovorum* subsp. *carotovorum* measured as change in absorbance at OD_600 nm_. **(B)** Protective effect of the phage cocktails on potato tuber slices against *Pectobacterium carotovorum* subsp. *carotovorum*. **(C)** Mean diameter of rotting tissue and **(D)** percent tissue maceration reduction in potato slices co-inoculated with *Pectobacterium carotovorum* subsp. *carotovorum* and phage cocktails.

#### 3.1.6. Phage-Mediated Biocontrol Using Monophage and Cocktail Treatments

##### 3.1.6.1. Monophage and cocktail treatments reduced soft rot symptoms in tissue slices

The average diameter of rotting tissue was significantly lower in all monophage treatments than in the set-up inoculated with *P. carotovorum* subsp. *carotovorum* only (Figure 3B-D). The set-up co-inoculated with *P. carotovorum* subsp. *carotovorum* and vB_PcaP-A3 exhibited the lowest diameter of rotting tissue among the three treatments (Figure 3B-D). Phage vB_PcaP-A3 was able to reduce potato tuber tissue maceration by at least 60%, while phages vB_PcaM-D1 and vB_PcaM-J3 were able to reduce tissue maceration by at least 50% of that observed for the control potato slices inoculated with *P. carotovorum* subsp. *carotovorum* only (Figure 3B-D). All of the phage cocktail treatments completely prevented tissue maceration at low and high MOI (Figure 4B-D).

##### 3.1.6.2. Advanced phage introduction can protect tubers from soft rot

The potential use of phages as a post-harvest disease control strategy was assessed using potato tubers artificially inoculated with *P. carotovorum* subsp. *carotovorum* treated with monophage suspensions applied at different time points. Results of the whole tuber assay showed that in the API treatment, monophage suspensions of the three phages applied on to the wound up to 30 h before bacterial inoculation provided protection against soft rot disease development (Figure 3E). In the DPI treatment, a 3-h and 8-h delay in the application of monophage suspensions of vB_PcaM-J3, and vB_PcaP-A3 and vB_PcaM-D1, respectively, following bacterial inoculation resulted in disease development (Figure 3E).

## 4. Discussion

Phage-mediated biocontrol of soft rot disease is a promising and an up-and-coming technology that has been widely explored worldwide. To our knowledge, this is the first report on the isolation and characterization of bacteriophages infecting *P. carotovorum* subsp. *carotovorum* in the Philippines, which explores some aspects that need to be considered for potential formulation of a postharvest biocontrol product against soft rot disease including phage stability, bioactivity in tissues, and application timing.

Three lytic bacteriophages infecting *P. carotovorum* subsp. *carotovorum* designated as vB_PcaP-A3, vB_PcaM-D1, and vB_PcaM-J3, were isolated from environmental samples including rhizosphere soil, symptomatic tissues, and sewage water. TEM analysis of the three phages revealed that vB_PcaP-A3 has a podovirus morphology similar to *Pectobacterium* phage Arno160, *Pectobacterium* phage vB_PatP_CB4, and *Dickeya* phage Amaethon (Buttimer, Hendrix, et al., 2018; Shneider et al., 2020). Meanwhile, phages vB_PcaM-D1 and vB_PcaM-J3 exhibited myoviral morphology similar to *Ackermannviridae Dickeya* phage PP35, *Pectobacterium* phage PP101, and *Pectobacterium* phage vB_PcaM_CBB (Kabanova et al., 2019; Lukianova et al., 2020).

The latent period and burst size of phages vB_PcaP-A3, vB_PcaM-D1, and vB_PcaM-J3 were determined based on one-step growth curve experiments. The phages were further characterized, and the adsorption rate, optimal MOI, and stability on varying temperature and pH levels were also determined. Of the three phage isolates, vB_PcaP-A3 has the lowest optimal MOI of 0.01, a latent period of 10 min and a calculated burst size value of 142 PFU/cell. Phage vB_PcaM-D1 has an optimal MOI of 0.1, latent period of 10 min and a burst size of 151 PFU/cell. vB_PcaM-J3 has an optimal MOI of 10, longest latent period of 30 min, and the lowest calculated burst size of 42 PFU/cell. Since phages displaying shorter latent periods and higher burst sizes are reported to provide better efficiency than phages with longer latent periods in cultures where the density of the host is high, vB_PcaM-J3 showed less promising growth characteristics compared to vB_PcaP-A3 and vB_PcaM-D1 (Abedon et al., 2003). All three phages, however, showed strong adsorption to their host evidenced by the observed 90% phage adsorption within 5 min after incubation. When their stabilities on varying temperature and pH levels were tested, all three phages survived incubations at 30°C-50°C and at pH 3-9, but were inactivated at 60°C and at pH 12. These are important environmental factors in biological control applications. Since temperatures under field conditions do not exceed 50°C and pH levels usually range from pH 3.3-4.3, results indicate that vB_PcaP-A3, vB_PcaM-D1 and vB_PcaM-J3 can be suitable for possible field applications (Navarrete et al., 2017).

All three phages showed specificity to their isolation host, *P. carotovorum* subsp. *carotovorum*. Phages are generally known to have narrow host ranges, typically limited to strains within a certain species of their bacterial host (Ross et al., 2016). This can allow the formulation of phage cocktails targeting bacterial species of interest only, which may be a certain bacterial plant pathogen or a specific bacterium in a microbial community that negatively impacts plant growth or health conditions. Developing a phage-based biocontrol product that can eliminate multiple genera, or even different related species could be challenging, but combining phages specific to target hosts may overcome this problem (Buttimer et al., 2017; Frampton et al., 2012). The specificity of phages also reduces the risk of harming the natural microbiota, especially those non-target bacteria that are potentially beneficial, and it limits the emergence of phage-resistant bacteria (Loc-Carrillo & Abedon, 2011).

In recent years, there were numerous attempts to systematize the single and cocktail application of phages to tubers and vegetable crops. For example, the first bacteriophage of *P. carotovorum* subsp. *carotovorum* that was sequenced, PP1, exhibited effective protection against its host in lettuce (Lim et al., 2013). In another study, two other phages namely □PD10.3 and □PD23.1 were applied together in both whole potato tubers and slices against a number of *Pectobacterium* and *Dickeya* isolates. Tissue maceration was significantly reduced by 80% and 95% in slices and whole tubers, respectively (Czajkowski et al., 2015). Two further studies revealed effective inhibition of SRPs in potato tubers using monophage formulations (J. Lee et al., 2017; Smolarska et al., 2018). In this study, the efficacy of the three bacteriophages, vB_PcaP-A3, vB_PcaM-D1, and vB_PcaM-J3, in both monophage and cocktail suspensions (combinations of two and three phages) were assessed through *in vitro* and semi-*in plant*a bacterial challenge tests, and we found that both monophage and phage cocktails completely inactivated the growth of *P. carotovorum* subsp. *carotovorum*. However, phages in cocktail suspensions were more effective over individual application in terms of their protective effect against soft rot by *P. carotovorum* subsp. *carotovorum* as indicated by the complete maceration reduction in both MOI (0.001 and 100) tested in the study. Recent reports of phage cocktail formulations against *P. carotovorum* subsp. *carotovorum* have also described efficient inhibition of bacterial growth both *in vitro* and *in vivo* (Carstens et al., 2019; Muturi et al., 2019; Zaczek-Moczydłowska et al., 2020). The use of higher MOI in phage formulations has also been reported to increase the chance for the bacteriophages to infect all host cells, thus effectively killing all host cells after one round of replication (Alves et al., 2016; Thomas, 2001).

Using different novel phage isolates against a number of SRP model strains, our work and the several studies mentioned above have provided preliminary evidence of the biocontrol potential of bacteriophages in controlling bacterial soft rot. However, a comprehensive understanding of their bioefficacy in postharvest applications is still limited and our work contributes to this pool of information by determining the efficient timing of phage treatment on tissues.

Preliminary experiments conducted to assess possible timing strategies in employing phage treatments to control post-harvest soft rot revealed the protective effect of introducing phages prior to bacterial inoculation (Figure 3E). In each monophage treatment, advanced phage introduction (API) treatments of up to 30 h were observed to protect the tubers from developing the disease (Figure 3E). With API, we hypothesize that, considering that the phage number applied is higher than or equal to the optimal MOI, the phages were able to rapidly kill the inoculated bacterial population, thus suppressing both the bacterial growth and the emergence of visible symptoms of the disease.

In contrast, a shorter window of time wherein the phage treatment seemed to offer effective protection from the disease was observed in the delayed phage introduction (DPI) treatment. This observation, in line with the hypothesized phage-bacterium interaction in the API treatment, seemed to suggest that a critical bacterial population relative to the phage population was required to effect noticeable disease manifestation. It was previously reported that quorum sensing regulates the coordination of the synthesis of virulence factors, including the production of plant cell wall degrading enzymes by *Pectobacterium* spp, which links the observed longer protective window of API treatment and the reduced protective window observed in DPI treatment (Pollumaa et al., 2012).

Furthermore, taking both into account the differences in the characteristics of the phages and the MOI used in this experiment (MOI 10.0, relative to bacterial population at time 0), it follows that, for the DPI treatment, a shorter protective window shall be observed for vB_PcaM-J3, which has the longest latent period (30 min), lowest burst size (42 CFU/cell) and highest MOI (10.0). That is, with the advanced introduction of the bacterial population of at least 3 h, the bacterial population seemed to have grown enough such that the vB_PcaM-J3 population is not at its optimal MOI anymore. In line with this is the observed longer protective window afforded by vB_PcaP-A3 and vB_PcaM-D1 that have lower optimal MOIs of 0.01 and 1.0 respectively, which are both lower than the MOI used in this experiment, and higher burst sizes of 151 CFU/cell and 142 CFU/cell, respectively. These characteristics may have also provided a buffering effect to account for the bacterial growth in DPI treatment.

Taken together, these results are suggestive of a potential post-harvest soft rot disease control strategy that may maximize the benefit of phage treatment by introducing efficiently lytic phages (ie. short latent period, larger burst size, and low optimal MOI) at a high MOI at the earliest time point after harvest. By doing so, while a small population of the soft rot-causing bacteria may have been inadvertently introduced during the process of harvesting, this population can be effectively controlled thus preventing bacterial growth, disease manifestation, and cross-contamination.

However, we are also aware that the use of higher MOI could also possibly increase the emergence of phage-resistant bacterial variants (Chen et al., 2018). This is due to the rapid evolution of the bacterial population from the dominance of phage-sensitive clones to phage-resistant clones when exposed to the selection pressure of lytic phages (Castillo et al., 2015; Hyman & Abedon, 2010; Middelboe, 2000; Middelboe et al., 2001). This implies that the use of heavy phage loads is an important selection factor that should be taken into account in phage therapy and that lower MOIs should also be used when testing protective effect and bacterial inactivation by phages, particularly in actual plant tissues (Chen et al., 2018). To this end, further experimental studies to optimize the phage treatment parameters and to identify the minimum effective phage number for potential field and post-harvest application are to be conducted.

## Data statement

All datasets presented in this paper are included in the article.

## CRediT authorship contribution statement

**Aeron Jade S. Parena:** Methodology, Validation, Formal Analysis, Investigation, Data Curation, Writing-Original Draft, Writing-Review & Editing, Visualization. **Benji Brayan I. Silva:** Conceptualization, Methodology, Validation, Formal Analysis, Writing-Review & Editing. **Rae Mark L. Mercado:** Methodology, Validation, Investigation, Data Curation. **Adrian Adelbert A. Sendon:** Methodology, Validation, Investigation. **Freddiewebb B. Signabon:** Methodology, Validation, Investigation. **Johnny F. Balidion:** Conceptualization, Supervision, Project Administration, Funding Acquisition. **Jaymee R. Encabo:** Conceptualization, Data Curation, Writing-Review & Editing, Supervision, Project administration, Funding acquisition.

## Acknowledgements

We thank Ms. Janren Sarah T. Macaraig and Mr. Benjamin V. Cunanan for the technical assistance with phage lysate preparations, and Mr. Aristeo P. Averion and Envirokonsult Equipment and Services, Inc. for the sewage water samples used for bacteriophage isolation.

