## Supplemental Table 1 for "Lytic phages displayed protective effects against soft rot-causing *Pectobacterium* sp."

**Supplementary Table 1.** Bacterial strains used in the study.

| **Species** | **Isolates** |
| --- | --- |
| *Pectobacterium carotovorum* subsp. *carotovorum* | BIOTECH1752 |
| *Pectobacterium aroidearum* | 18a |
| *Pectobacterium aroidearum* | 2Cab 3_2 C15 |
| *Pectobacterium aroidearum* | 109L |
| *Pectobacterium aroidearum* | 120b |
| *Pectobacterium aroidearum* | 1BCara |
| *Pectobacterium aroidearum* | 1Cab9a |
| *Pectobacterium aroidearum* | 1Cab15a |
| *Pectobacterium aroidearum* | 2Cab15b |
| *Pectobacterium aroidearum* | 1Car4 |
| *Pectobacterium aroidearum* | 2Car3 |
| *Pectobacterium aroidearum* | 2Car10B |
| *Pectobacterium aroidearum* | 1EPbb2 |
| *Pectobacterium aroidearum* | 2P2 |
| *Pectobacterium aroidearum* | 1Pb |
| *Pectobacterium aroidearum* | 2Pb3P |
| *Pectobacterium aroidearum* | 37a |
| *Pectobacterium aroidearum* | *39a* |
| *Pectobacterium aroidearum* | 3a |
| *Pectobacterium aroidearum* | 46 |
| *Pectobacterium aroidearum* | 56b |
| *Pectobacterium aroidearum* | *57a* |
| *Pectobacterium aroidearum* | 58a |
| *Pectobacterium aroidearum* | 5L |
| *Pectobacterium aroidearum* | 68b |
| *Pectobacterium aroidearum* | 71/C4 |
| *Pectobacterium aroidearum* | 72 |
| *Pectobacterium aroidearum* | 73 |
| *Pectobacterium aroidearum* | 74a |
| *Pectobacterium aroidearum* | 75a |
| *Pectobacterium aroidearum* | 78a |
| *Pectobacterium aroidearum* | 7V |
| *Pectobacterium aroidearum* | 7V |
| *Pectobacterium aroidearum* | 83a |
| *Pectobacterium aroidearum* | 84b |
| *Pectobacterium aroidearum* | 89a |
| *Pectobacterium aroidearum* | 8EP2 |
| *Pectobacterium aroidearum* | MO2 |
| *Pectobacterium aroidearum* | MO21a |
| *Pectobacterium aroidearum* | MO52 |
| *Pectobacterium aroidearum* | PcC |
| *Pectobacterium cacticida* | MO46a |
| *Pectobacterium carotovorum* | 2Cab13b |
| *Pectobacterium carotovorum subsp. brasiliense* | 11cc |
| *Pectobacterium carotovorum subsp. brasiliense* | 28cc |
| *Pectobacterium carotovorum subsp. brasiliense* | 90c |
| *Pectobacterium carotovorum* | 2ACar |
| *Pectobacterium odoriferum* | - |
| *Pectobacterium wasabiae* | 15a |
| *Escherichia coli* | - |
| *Bacillus subtilis* | - |
| *Staphylococcus aureus* | - |
| *Pseudomonas aeruginosa* | - |
| *Pseudomonas putida* | - |
| *Proteus vulgaris* | - |
| *Salmonella enterica* | JCM1651 |
| *Xanthomonas oryzicola* | - |
